# Partial refuges from biological control due to intraspecific variation in protective host traits: a case study with the egg parasitoid *Trissolcus japonicus*

**DOI:** 10.1101/2022.04.28.489927

**Authors:** Paul K. Abram, Tim Haye, Peggy Clarke, Emily Grove, Jason Thiessen, Tara D. Gariepy

**Affiliations:** Agriculture and Agri-Food Canada, Agassiz Research and Development Centre, Agassiz, BC, Canada; CABI, Delémont, Switzerland; Simon Fraser University, Department of Biological Sciences, Burnaby, BC, Canada; Agriculture and Agri-Food Canada, London Research and Development Centre, London, ON, Canada

**Keywords:** Intraspecific variation, non-target effects, refuge theory, *Halyomorpha halys*, *Trissolcus japonicus*

## Abstract

Predicting how much of a host or prey population may be attacked by their natural enemies is fundamental to several subfields of applied ecology, particularly biological control of pest organisms. Hosts or prey can occupy refuges from natural enemy attack, but habitat or ecological refuges are challenging or impossible to predict in a laboratory setting – which is often where efficacy and specificity testing of candidate biological control agents is done. Here we explore how intraspecific variation in continuous traits that confers some protection from natural enemy attack – even after the natural enemy has encountered the prey – could provide partial refuges. The size of these trait-based refuges should depend on the relationship between trait values and host/prey susceptibility to natural enemy attack, and on how common different trait values are within a host/prey population. These can be readily estimated in laboratory testing of natural enemy impact on target or non-target prey or hosts as long as sufficient host material is available. We provide a general framework for how intraspecific variation in protective host traits could be integrated into biological control research, specifically with reference to non-target testing as part of classical biological control programs. As a case study, we exposed different host clutch sizes of target (pest) and non-target (native species) stink bug (Hemiptera: Pentatomidae) species to a well-studied exotic biocontrol agent, the egg parasitoid *Trissolcus japonicus* (Hymenoptera: Scelionidae). Although we observed several behavioural and reproductive responses to variation in host egg mass size by *T. japonicus*, they did not translate to increases in predicted refuge size (proportion host survival) large enough to change the conclusions of non-target testing. We encourage researchers to investigate intraspecific variation in a wider variety of protective host and prey traits and their consequences for refuge size.

## Introduction

Estimating the proportion of host or prey populations attacked by a natural enemy is a foundational part of estimating the efficacy and environmental safety of biological control agents across a wide variety of contexts. As one example, during routine host specificity testing of candidate biological control agents for invasive pests, ‘non-target’ species (i.e., potential hosts or prey not being targeted by the biological control agent) are exposed to candidate agents in laboratory microcosm tests to determine whether they will attack species other than the target pest (Van Driesche and Reardon 2004). While these fundamental (or physiological) host range trials provide important information to reduce the likelihood of unintended negative ecological impacts (van Lenteren et al. 2006), these tests are highly conservative and probably overestimate the number of host species attacked under field conditions (Haye et al. 2005; Barratt et al. 1997, 2010; Cameron et al. 2013). In addition, while they can predict what host or prey species *could* be attacked by agent, they provide poor estimates of whether or not the natural enemy would significantly suppress non-target host populations, and if so by how much (Louda et al. 2003). This is an important distinction, as in some contexts a certain level of non-target host use by released agents could be considered acceptable – especially if it does not lead to significant suppression or extirpation of populations of non-target species (Van Driesche and Hoddle 2017). Identify factors that reduce the susceptibility of non-target species to attack by biological control agents would be part of a recent trend towards incorporating more ecological realism in laboratory host range methodology to improve their predictive power (Haye et al. 2020).

Population-level impact of biological control agents can be mitigated if some proportion of the host or prey population is not susceptible to attack by the biological control agent – that is, they occupy a refuge (Hawkins et al. 1993). Refuges from attack by natural enemies can occur at different spatial and temporal scales and due to a variety of mechanisms (Gross 1993; Mills and Getz 1996). For example, hosts or prey may occupy a habitat, feeding niche, or host plant where levels of attack by a natural enemy are lower (Johnson et al. 2005), or may be protected from natural enemies by mutualistic ecological interactions (Wyckhuys et al. 2007). These refuges may be due to pre-existing niche preferences (Johnson et al. 2005; Wyckhuys et al. 2007), or can shift due to new interactions with natural enemies (e.g., Murphy 2004). Hosts or prey may also pass through life stages that are less susceptible to natural enemy attack (e.g. living in concealed feeding niches), disperse to new areas before their natural enemies, reproduce during times of the season when the natural enemy is not present, or may only be attacked by the natural enemy during relatively brief ‘spillover’ events (Mills and Getz 1996; Strom et al. 2001; Catton et al. 2015). What these spatial and temporal ‘ecological’ refuges have in common is that they lower the proportion of hosts that natural enemies encounter and thus have the opportunity to attack. However, some individuals of host or prey populations, even in their most susceptible life stages, may occupy refuges because they possess protective characteristics that make them more likely to survive encounters with natural enemies; these include physiological defenses (e.g., immunity, chemical defenses), morphological traits (e.g., chorion/integument/puparia thickness, hairs, spines, silk cocoons), behavioural defenses (e.g., kicking, dropping), and forming large aggregations that saturate natural enemies (reviewed in Gross 1993). The total size of a refuge occupied by a given host or prey population would be the sum of the proportions occupying ‘ecological’ refuges (i.e., what proportion are not found by their natural enemy) and ‘trait-based’ refuges (i.e., what proportion found by natural enemies survive the encounters).

The amount of protection conferred by a continuous trait often depends on the trait’s value, which would usually vary among host or prey individuals. There have been several documented cases of variation in protective traits within and among host and prey populations (Cronin and Gill 1989; Henter 1995; Kraaijeveld and van Alphen 1999; Abrams 2000 and references therein). The size of a trait-based refuge can be described by the product of two functions: the frequency distribution of the trait’s value in the host or prey population (i.e. how common each trait value is in the population), and the relationship between the trait’s value and the amount of protection conferred against natural enemy attacks (Figure 1). We suggest that although spatial and temporal ecological refuges would be hard (if not impossible) to identify in laboratory host range testing, many hypothesized trait-based refuges could be relatively easily measured and accounted for. Traits of hypothesized importance in non-target species subjected to host range testing could be manipulated experimentally or simply measured and included as covariates in analyses of host or prey susceptibility to attack by candidate biocontrol agents. In addition to identifying potential trait-based refuges within the species tested, explicitly considering how intraspecific trait variation in hosts relates to susceptibility to a given natural enemy could hypothetically provide clues to the susceptibility of untested species who also possess variation in that trait. Despite much interest in the ecological importance of intraspecific variation in fundamental ecological research (Bolnick et al. 2011; Des Roches et al. 2018), non-target studies, as far as we are aware, still focus mostly on interspecific trait differences, and either hold host phenotypes as constant as possible or do not explicitly account for their variation (e.g., Table 1). In biological control of plants, which are well-known to exhibit intraspecific variation in defensive chemistry (Hahn and Maron 2016), consideration of intraspecific variation in phenotypic traits is more common (e.g., Paterson et al. 2012; Stutz et al. 2021; Sun et al. 2022), although we are not aware of cases in biological control where it has been considered for quantitatively estimating the size of refuges from natural enemy attack.

**Figure 1.**
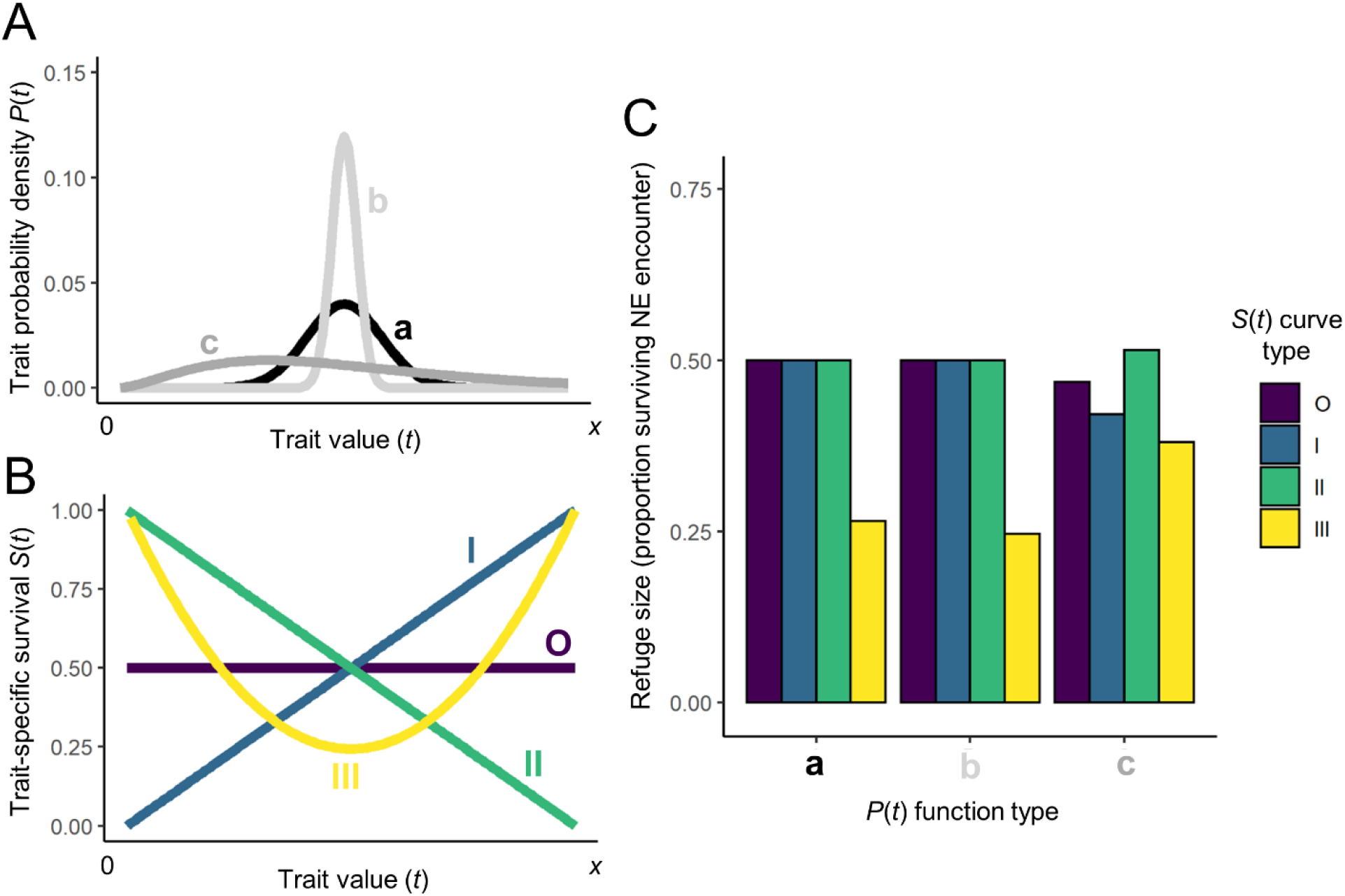
The size of a refuge due to intraspecific variation in a protective trait is the product of the proportion of individuals with each value of the trait, and the amount of protection conferred by each trait value. (A) The value of a trait, *x*, follows one of three probability density relationships, *P*(*t*), with the same mean but different distributions: (a) a normal distribution with mean *j* and standard deviation *k*; (b) a normal distribution with mean *j* and standard deviation *k*/3; (c) a binomial distribution with mean *j* and dispersion parameter *j*/20. (B) The effect of the trait’s value on susceptibility to parasitism. The different trait-specific survival functions, *S*(*t*) all have the same mean value (0.5) over the range of values 0→*x*. (C) The proportion of a host or prey species’ population occupying a refuge from natural enemy (NE) attack after encounter, calculated for each combination of the relationships in panels (A) and (B), as 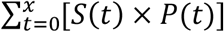. The different combinations of trait densities and survival curves, despite each having the same mean values, can yield different total proportions of surviving hosts depending on the characteristics of these two functions. Note, for example, that asymmetric *P*(*t*) (e.g. ‘c’) and/or non-linear *S*(*t*) (e.g., III) can generate different refuge sizes for the same mean level of susceptibility across all values of *t*.

**Table 1.** Previous no-choice studies testing the host range of *Trissolcus japonicus*, with: the number of non-target (NT) Hemiptera species found to be susceptible (i.e., *T. japonicus* was shown to reproduce on and/or kill the species’ eggs) and how many were tested (in addition to *Halyomorpha halys* acting as a control, unless otherwise indicated); the number of egg masses tested per NT species; whether the NT egg mass sizes were unmanipulated (i.e. naturally occurring sizes from insect colonies were used), held constant (manipulated to be the same or similar [∼] size in every trial), or intentionally manipulated (intentionally made larger and smaller than naturally occurring egg masses, but encompassing the natural range); whether direct behavioural observations of parasitoids were conducted with videos or live observations; whether non-reproductive effects (NREs) of parasitoids on hosts were estimated by comparing mortality of exposed egg masses to controls; and arena type in which parasitoids were exposed to non-target hosts.

| Study citation | Region | Number of NT species susceptible/tested | Number of egg masses tested per NT species (range) | Treatment of NT egg mass sizes | Direct behavioral observations | Measurement of NREs with controls | Arena type |
| --- | --- | --- | --- | --- | --- | --- | --- |
| Zhang et al. (2017) | China | 7/8 | 4 to 61 | Unmanipulated | No | No | Petri dish |
| Hedstrom et al. (2017) | USA | 7/10 | 19 to 27 | Unmanipulated | No | No | Vial |
| Haye et al. (2019) | Switzerland/Italy | 12/13 | 16 to 46 | Constant (10 eggs)<sup>a</sup> | Yes | Yes | Petri dish |
| Lara et al. (2019) | USA | 6/10 | 8 to 50 | Unmanipulated<sup>a</sup> | No | No | Vial |
| Charles et al. (2019) | New Zealand | 6/7<sup>b</sup> | 19 to 70 | Unmanipulated | No | No | Vial |
| Boyle et al. (2020) | USA | 1/1 | 25 | Constant (~27 eggs) | No | No | Cage with plant |
| Sabbatini-Peverieri et al. (2021) | Italy | 8/15 | 8 to 43 | Constant (1 egg) | Yes<sup>c</sup> | Yes | Vial |
| Saunders et al. (2021) | New Zealand | 1/1<sup>b</sup> | 15 | Unmanipulated | Yes | Yes<sup>d</sup> | Vial |
| This study | Canada | 3/3 | 40 to 44 | Manipulated | Yes | Yes | Mesocosm with plant |

<sup>a</sup> In these studies, there were exceptions to whether egg masses were manipulated or not for single host species
<sup>b</sup> Did not include *H. halys* in testing as a control
<sup>c</sup> Conducted behavioural observations for first 2h of 24h exposure period
<sup>d</sup> Unexposed egg masses were 'pseudo-controls' collected during a different time period from host range trials.

*Trissolcus japonicus* (Ashmead) (Hymneoptera: Scelionidae) is a biological control agent of the invasive brown marmorated stink bug, *Halyomorpha halys* (Stål) (Hemiptera: Pentatomidae), with both species originating in Asia (Hoebeke and Carter 2003; Abram et al. 2020; Conti et al. 2021). *Trissolcus japonicus* is currently being intentionally released or redistributed in its adventive range in Europe and North America (Abram et al. 2020; Conti et al. 2021). Extensive host specificity testing with non-target true bug (Hemiptera) species has been done worldwide to inform these biological control programs (Table 1). Although a number of non-target species are susceptible to attack by *T. japonicus*, interpretation of some results is hindered by the fact that parasitoid behaviour was not observed, or unexposed control egg masses were not included in experimental designs to estimate how many hosts were killed without parasitoid reproduction (i.e., non-reproductive effects; Abram et al. 2016; Abram et al. 2019). While *T. japonicus* is known to attack some non-target stink bugs (Milnes and Beers 2019; Hepler et al. 2020), the population-level impact of this attack is not known (Kaser et al. 2018; Abram et al. 2020). Ecological refuges from parasitism (e.g. habitats at higher altitude) are thought to have mitigated population-level impacts of a closely related egg parasitoid, *Trissolcus basalis* (Wollaston) on a native Hawaiian stink bug species (Johnson et al. 2005). The potential for trait-based non-target host refuges from parasitoid attack merits investigation as well.

In this study, we tested whether intraspecific variation in the size of egg masses of target and non-target hosts of *T. japonicus* could theoretically provide partial refuges from parasitism. Stink bugs lay their eggs in aggregated egg masses that vary inter- and intra-specifically in clutch size (the number of eggs in each egg mass) (Abram et al., in prep.). While clutch size is not technically a trait of each individual egg, per se, it essentially functions as such, as it can determine the per-capita survival of eggs within a clutch and is thus subject to selective pressures that cause individual egg mortality (Godfray et al. 1991). None of the aforementioned host range testing studies have explicitly considered the effect of host egg mass size on the observed variation in host susceptibility to *T. japonicus*: host egg mass size was either held constant at sometimes unrealistic egg mass sizes, or unmanipulated egg masses were used but the effect of size variation was not quantified (Table 1).

We hypothesized that egg mass size could be an important aspect of host quality for ‘quasi-gregarious’ parasitoids such as *T. japonicus* for which an egg mass essentially represents a single host, and the number of eggs in a clutch determines how many offspring they will produce. Specifically, we predicted that eggs in the smallest and largest egg masses may be at less risk of mortality when encountered by *T. japonicus*. Parasitoids are known to be more likely to reject relatively low-quality (including smaller-sized) hosts in favour of higher quality hosts (e.g., Heimpel et al. 1996; reviewed in McGregor and Roitberg 2000); thus, we predicted that *T. japonicus* may choose to reject smaller egg masses, especially given that its most closely-associated host, *H. halys*, lays relatively large egg masses (usually 28 eggs). In addition, we predicted that some eggs in larger egg masses could be more likely to survive if the egg mass size is large enough to cause *T. japonicus* to deplete their egg loads, as has been shown for other parasitoids (Weseloh 1972; Laumann et al 2008; in Mills and Getz 1996 and references therein). We predicted that refuges from parasitism at small and large egg mass sizes would favor survival of stink bug eggs at the relative extremes of the host species’ clutch size distributions, resulting in a ‘type III’ trait-specific survival relationship as depicted in Figure 1. We further hypothesized that this relationship would interact with the distribution of egg mass sizes laid by stink bug populations to determine the proportion of egg masses occupying a refuge from parasitism. For example, the type III trait-specific survival relationship would be predicted to generate larger refuges for stink bug species that lay more variably-sized egg masses (Figure 1). This study is intended as an illustrative first step in considering how intraspecific variation in host traits could affect the interpretation of host specificity testing in biological control programs.

## Material and Methods

### Insect rearing

The exotic *Halyomorpha halys* and three ‘non-target’ North American stink bug (Hemiptera: Pentatomidae) species, *Podisus maculiventris* (Say), *Euschistus conspersus* Uhler, and *Chlorochroa ligata* (Say), were tested based on availability and ease of rearing, and because they span a range of lifestyles, egg mass sizes, and known levels of suitability as hosts for *T. japonicus* (references in Table 1). Further, the distribution of egg mass sizes (i.e., number of eggs per clutch laid), as well as the characteristics of their frequency distributions (skewness, amount of variation around the mean) varies considerably among the four stink bug species in this study (Figure 2).

**Figure 2.**
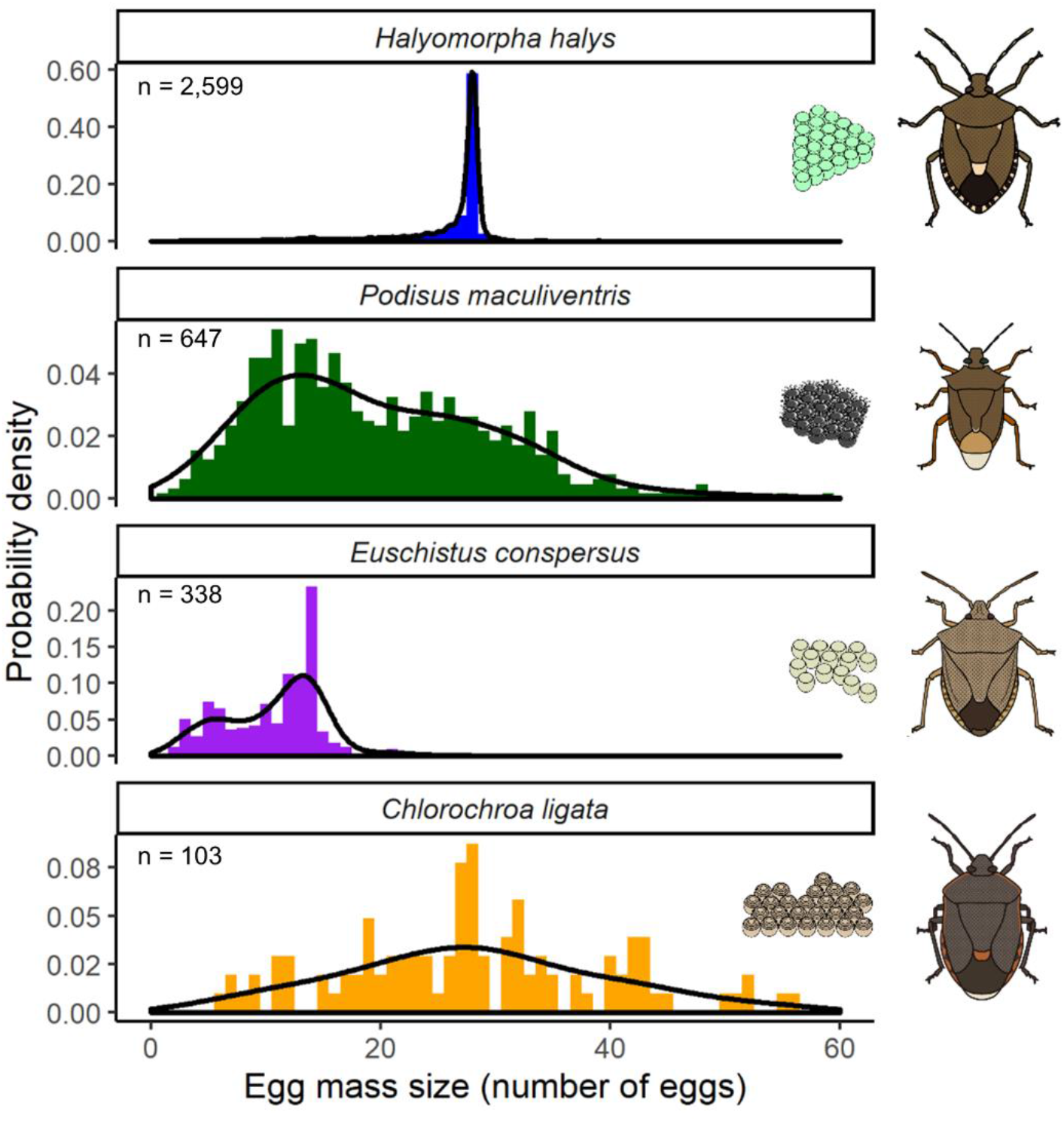
Egg mass size probability density distributions for the four stink bug species whose eggs were exposed to *Trissolcus japonicus* in this study. Data is from multiple pooled laboratory and field datasets from different research groups, compiled in the clutch size database in Abram et al. (in prep.). Black curves are probability density functions using a Gaussian smoothing kernel with a smoothing bandwidth equal to 1.4 times the standard deviation of the smoothing kernel (set arbitrarily to maximize visual fit to distributions). Note the differences in the y-axis scales.

*Halyomorpha halys, C. ligata*, and *E. conspersus* were collected from field populations in the Fraser Valley region of British Columbia, Canada in 2018 and 2019 and reared on mixed diets of potted plants, seeds, and fruit as described in Wong et al. (2021). *Podisus maculiventris* was sourced from collections in Ontario, Canada and reared on larvae and pupae of *Tenebrio molitor* L., and potted fava bean plants as a source of water and supplemental nutrition. Adults stink bugs were kept in large mesh cages (45 × 45 × 45 cm; Megaview, Taiwan) at 23 ± 2ºC, 40-55% RH, and a 16:8 h L:D photoperiod with Reemay polyester fabric (Avintiv, USA) as the primary oviposition substrate. Egg masses were collected every 1-2 days and were <48h old when used in trials.

The *T. japonicus* laboratory colony was established from individuals originally collected in Beijing, China in 2009 by the United States Department of Agriculture, Agricultural Research Service (USDA-ARS). This is the same *T. japonicus* colony used in host range testing studies in North America, New Zealand, and Europe (Hedstrom et al. 2017; Charles et al. 2019; Lara et al. 2019; Boyle et al. 2020; Saunders et al. 2021; Sabbatini-Peverieri et al. 2021; Table 1). Parasitoids were maintained on newly laid (< 48h) eggs of *H. halys* in ventilated plastic containers (width: 9.2 cm; height: 11.7 cm) and reared at 23 ± 1ºC, 25-50% RH, and a 16:8 h L:D photoperiod. Femaleswere 4-10 days old when used in experiments to allow time for ovary maturation (Wong et al. 2021; R. Paul, J. Lee and P.K. Abram, accepted). Female *T. japonicus* used in experiments emerged in the presence of males, and thus were assumed mated. They were isolated in 1.7 mL plastic microcentrifuge tubes with a drop of liquid honey within 48h of emergence until released into experimental arenas.

### Egg mass size manipulation

To experimentally test the effect of different egg mass sizes on susceptibility of stink bug egg masses to *T. japonicus* with adequate replication across a large variety of egg mass sizes, a range of egg mass sizes was created for each species by sequentially assigning egg masses, as they were collected, to one of the following four conditions: (i) normal-sized (the number of eggs were not manipulated); (ii) double the normal size (two egg masses were placed next to each other); (iii) half of the normal size (half of the eggs in each egg mass were removed); and (iv) a quarter of the normal size (three quarters of eggs in each egg mass were removed). Because there is variation in the size of unmanipulated egg masses (Figure 2), these four categorical treatments created a variety of egg mass sizes for each species, and egg mass size was treated as a continuous factor in subsequent analyses. Resultant egg mass sizes varied within species as follows: *H. halys*: 5–59; *P. maculiventris*: 2–41, *E. conspersus*: 3–31; *C. ligata*: 5–81. These ranges span a range of minimum and maximum values similar to that of unmanipulated egg mass sizes for each species (Figure 2). Mean sizes of unmanipulated egg masses for each species were as follows: *H. halys*: 27.3 ± 0.2; *P. maculiventris*: 16.2 ± 1.5, *E. conspersus*: 14.3 ± 0.3; *C. ligata*: 28.6 ± 3.4 (means ± SEs). To measure baseline egg mortality in the absence of parasitism, and ensure that egg mortality was not associated with egg mass size (or egg mass size manipulation), masses from each size category were assigned to one of two conditions: (A) not exposed to parasitoids; (B) exposed to parasitoids (see below). Egg masses not exposed to parasitoids were collected within the same week as paired exposed egg masses and were treated the same way in terms of how their sizes were manipulated (or not), what substrate they were fixed to, exposure to adhesive, and incubation conditions (see below). We set aside size category-matched unexposed egg masses in parallel when exposing egg masses to parasitism whenever possible, but during some time periods the number of available egg masses was limited and so exposed egg masses were prioritized. Trials for different non-target species (*P. maculiventris, E. conspersus, C. ligata*) were run during different time periods due to different seasonal availability of egg masses, but we ran simultaneous matched trials with *H. halys* eggs on every day that experiments were conducted. After verifying that the survival of unexposed *H. halys* eggs and the suitability of *H. halys* eggs for *T. japonicus* were similar over time, we pooled the trials with *H. halys* for analysis, and analyzed the results for each of the four stink bug species separately (see below). Total sample sizes for exposed and unexposed egg masses were as follows: *H. halys*: 141 exposed, 146 unexposed; *P. maculiventris*: 40 exposed, 40 unexposed; *E. conspersus*; 40 exposed, 36 unexposed; *C. ligata*: 44 exposed, 35 unexposed.

### Experimental setup

Experiments where egg masses of different sizes were exposed to *T. japonicus* were conducted under the same abiotic conditions as those under which the parasitoids were reared (see above). Arenas where trials took place consisted of 1 litre cylindrical plastic containers containing a soybean (*Glycine max* [L.]) seedling with 4-6 true leaves (Appendix S1: Figure S1). There were eight of these arenas set up adjacent to each other, allowing trials for different egg mass size categories to be run for one non-target stink bug species as well as *H. halys* simultaneously.

Arenas were designed to be large enough and contain enough structural complexity to allow for more realistic parasitoid foraging and patch exploitation behaviour than in most past studies of *T. japonicus* host range (Table 1). On the morning of the day they were to be exposed to parasitoids, egg masses fixed on squares of polyester fabric were glued to the underside of one of the true leaves of the soybean plant with white non-toxic glue (School Glue, LePage, Canada).

Containers with the plants and egg masses were set on an elevated transparent plexiglass surface, allowing continuous filming of the egg mass from below with video camera (DinoLite, Canada). Parasitoids were released into arenas between 7 and 9 hours after the beginning of the photophase, and the egg mass in each arena filmed until 2 to 3 hours after the start of photophase the following morning to record parasitoid foraging behaviour (=17-20 hours of filming per trial). The 7-9 hours of filming during the first day occurred during the peak activity times of *T. japonicus* (R. Paul, P.K. Abram, and J. Lee, accepted) and was based on preliminary trials to measure host finding success and parasitism rate, and was designed to give parasitoids more than adequate time to find and parasitize all eggs that their egg loads would allow, even in the largest egg masses. Egg masses were then removed from arenas and incubated under the same abiotic conditions for at least 40 days to record either stink bug nymph or parasitoid emergence. After insect emergence concluded, we dissected remaining eggs to count the number of unemerged stink bug nymphs, unemerged parasitoid adults or pupae, and aborted eggs with undiscernible contents. Eggs were then classified as either yielding an emerging adult parasitoid, emerged stink bug nymphs (i.e., surviving hosts), or aborted, where both the host and parasitoid die, as per the definition of Abram et al. (2016) (containing parasitoid adults or pupae that failed to emerge or undiscernible contents, which could include parasitoids that died in the egg or larval stage). Host eggs that aborted for reasons other than unsuccessful parasitism *versus* because of unsuccessful parasitism cannot be readily distinguished without molecular diagnostic tools (e.g. Hepler et al. 2020), but comparisons of abortion levels of eggs exposed to parasitoids with those in unexposed control egg masses can allow an estimate of how much host egg abortion is due to parasitoids (Abram et al. 2016).

### Video analysis of parasitoid behaviour

Videos were reviewed to record when (if ever) parasitoids first encountered the egg mass (i.e., contacted it with their legs and antennae), the number and temporal rank of ovipositions (ovipositior insertions + marking behaviour; see Abram et al. 2014) and ovipositor rejections (ovipositor insertion without marking behaviour) of host eggs, and when (if ever) the parasitoid first left the egg mass for more than 30 minutes (a semi-arbitrary threshold based on the observed distribution of leaving times in *T. japonicus*). The analysis focused on the first 7-9 hours of filming during the first day’s photophase, but the observers reviewed the 2-3 hours of video from the beginning of the next day’s photophase to verify that additional ovipositions or rejections did not occur during that time. We used these observations to determine parameters that would allow us to address the study’s main predictions about how parasitoid behaviour would respond to egg mass size. First, we created a binary (0, 1) variable to indicate whether parasitoids ever found (=contacted) each egg mass or not during the filming period. Next, to test the prediction that parasitoids would be more likely to accept larger egg masses, we created a binary (0, 1) variable, ‘acceptance probability’, recording whether or not parasitoids that encountered egg masses subsequently oviposited in at least one host egg. To test the prediction that parasitoids would exhaust their egg loads on larger egg masses, we used the number of observed ovipositions per available host egg as a proxy for egg load depletion, reasoning that for a given host species-specific level of acceptability, parasitoids would oviposit fewer times relative to the number of eggs in the egg mass when they had depleted their egg loads, with values close to 1 indicating that parasitoids had adequate egg loads to exploit every host egg and values less than 1 indicating that several hosts would remain unparasitized (values were sometimes more than 1 if parasitoids oviposited in individual host eggs multiple times). Additional behavioural analyses of oviposition rate, post-oviposition guarding behaviour, and probability of ovipositor rejections over time are described in Appendix S1: Methods S1.

### Statistical analysis

All statistical analyses were done with R version 3.6.1 (R Core Team 2019). Generalized linear models (GLMs) were used to determine the effect of egg mass size on each of the behavioural and developmental parameters for each stink bug host species (see Table 2). In the case of host survival, we also tested the statistical significance of a quadratic term (egg mass size^2^), given the relationship predicted by a type III survival curve (see Figure 1). The type of GLM (the error distribution and link function) depended on the type and distribution of the response variable (Table 2). The default link functions were used in all cases. The assumptions of each model were checked by inspecting quantile-quantile and residual plots of model fits, and examining model outputs for evidence of overdispersion; quasi-likelihood was used when model fits were overdispersed. Significance testing (p < 0.05) was done with likelihood ratio tests, except for models using Gaussian or quasi-binomial error distributions, in which case F-tests were used instead (Crawley 2012). The R packages we used for data analysis and visualization were ‘car’ (Fox and Weisberg 2019) and ‘ggplot2’ (Wickham 2016).

**Table 2.** Statistical significance of host egg mass size (EMS) as a predictor of behavioural and developmental parameters of *Trissolcus japonicus* exploiting the eggs of four different stink bug species. For the proportion of hosts surviving, statistics for quadratic terms are not reported when not statistically significant (p > 0.05). Statistics are bolded when EMS is a significant (p < 0.05) predictor of a given response variable.

| Response variable | Host species |  |  |  |
| --- | --- | --- | --- | --- |
|  | *H. halys* | *P. maculiventris* | *E. conspersus* | *C. ligata* |
| Probability of finding egg mass <sup>a</sup> | $\chi^2_{1,139} = 1.2$<br>$p = 0.27$ | $\chi^2_{1,38} = 0.9$<br>$p = 0.33$ | $\chi^2_{1,38} = 1.2$<br>$p = 0.27$ | $\chi^2_{1,42} = 1.2$<br>$p = 0.41$ |
| Patch acceptance probability <sup>a</sup> | $\chi^2_{1,104} = 2.29$<br>$p = 0.13$ | $\chi^2_{1,29} = \mathbf{5.9}$<br>$\mathbf{p = 0.015}$ | $\chi^2_{1,27} = 0.0$<br>$p = 1.00$ | $\chi^2_{1,34} = \mathbf{9.3}$<br>$\mathbf{p = 0.0023}$ |
| Ovipositions per host egg <sup>b</sup> | $\mathbf{F_{1,103} = 60.9}$<br>$\mathbf{p < 0.0001}$ | $F_{1,103} = 1.05$<br>$p = 0.31$ | $F_{1,25} = 1.0$<br>$p = 0.32$ | $\mathbf{F_{1,25} = 37.9}$<br>$\mathbf{p < 0.0001}$ |
| Proportion hosts surviving <sup>c</sup> | $F_{1,106} = 0.31$<br>$p = 0.58$ | $F_{1,28} = 2.6$<br>$p = 0.12$ | $F_{1,32} = 0.18$<br>$p = 0.68$ | $\mathbf{EMS: F_{1,33} = 9.8; p = 0.0036}$<br>$\mathbf{EMS^2: F_{1,33} = 10.1; p = 0.0033}$ |
| Wasp offspring per host egg <sup>c</sup> | $\mathbf{F_{1,101} = 48.3}$<br>$\mathbf{p < 0.0001}$ | $F_{1,25} = 0.5$<br>$p = 0.48$ | $\mathbf{F_{1,26} = 4.7}$<br>$\mathbf{p = 0.039}$ | $F_{1,25} = 0.2$<br>$p = 0.69$ |
| Proportion aborted host eggs <sup>c</sup> | $\mathbf{F_{1,101} = 20.4}$<br>$\mathbf{p < 0.0001}$ | $F_{1,25} = 0.01$<br>$p = 0.92$ | $F_{1,26} = 1.2$<br>$p = 0.28$ | $F_{1,25} = 3.1$<br>$p = 0.089$ |

<sup>a</sup> GLM with binomial error distribution
<sup>b</sup> GLM with Gaussian error distribution (=linear models)
<sup>c</sup> GLM with quasi-binomial error distribution

## Results

### Egg mass acceptance and number of ovipositions per host egg

*Trissolcus japonicus* females encountered egg masses of different sizes with equal probability, with between 77% and 92% of host egg masses depending on host species (Table 2). For egg masses encountered by *T. japonicus, P. maculiventris* and *C. ligata* egg masses with fewer eggs were less likely to be accepted, while acceptance probability was insensitive to egg mass size for

*E. conspersus* and *H. halys* (Table 2; Figure 3). For accepted egg masses, *T. japonicus* oviposted fewer times per available host egg in larger *H. halys* and *C. ligata* egg masses (also the two species with the largest egg masses overall), but oviposited a similar number of times per host egg in different egg mass sizes for the other two host species (Table 2; Figure 3).

**Figure 3.**
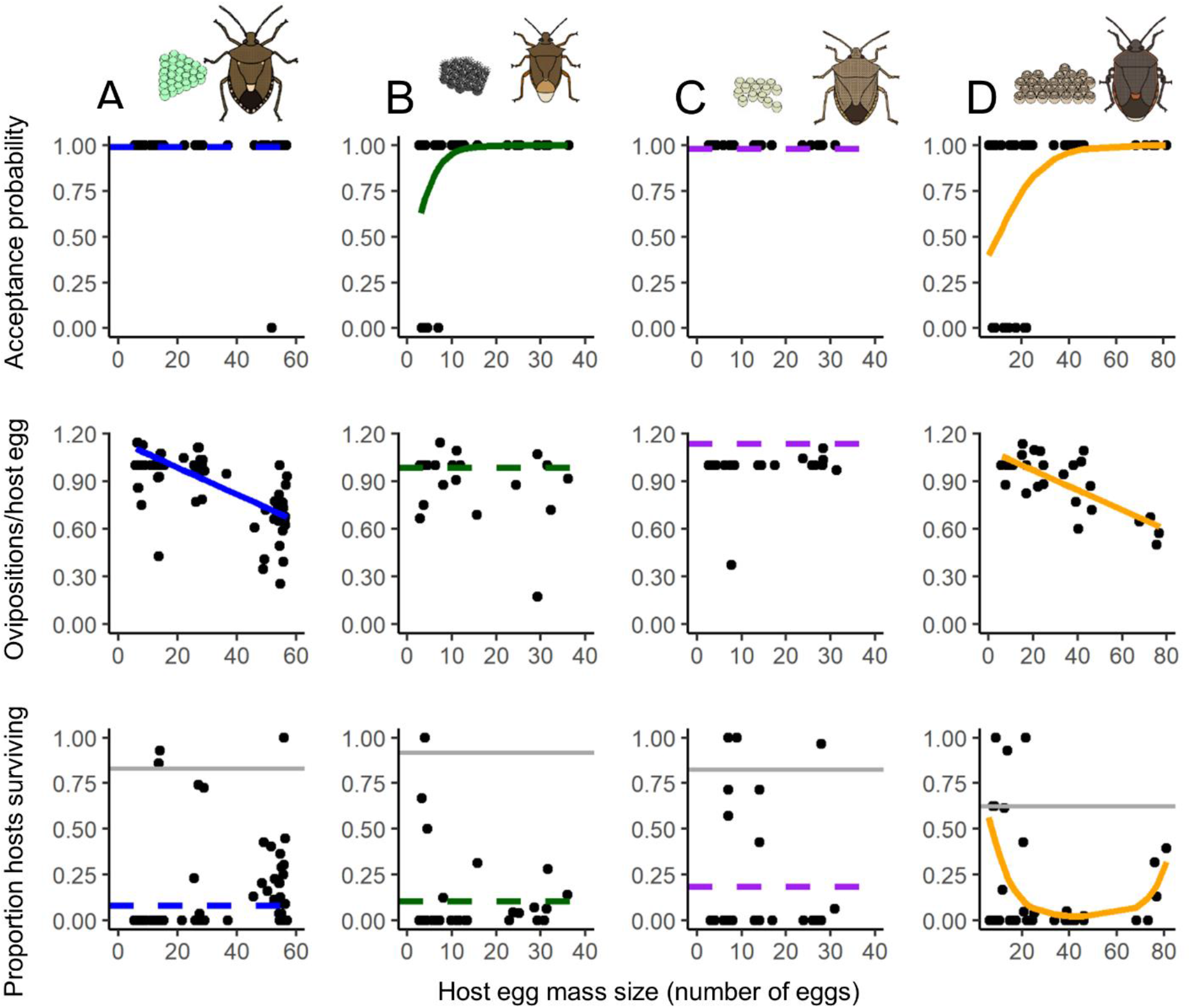
The effect of host egg mass size on *Trissolcus japonicus* patch acceptance probability (for wasps that found the egg mass, the probability that they will oviposit in at least one host egg after finding the host egg mass), the number of ovipositions per host egg (for wasps that found and accepted the egg mass), and the proportion of surviving hosts (for egg masses that were found by wasps; solid grey lines show mean host survival in egg masses not exposed to parasitoids) for eggs of four different host species: A) *Halyomorpha halys*; B) *Podisus maculiventris*; C) *Euschistus conspersus*; D) *Chlorochroa ligata*. Solid lines show statistically significant GLM fits; horizontal dashed lines show global means when the relationship between the response variable and host egg mass size was not statistically significant (see Table 2 for statistical information). Note that to keep vertical scales consistent, two high-value points are omitted from the “Ovipositions/host egg” plot in C).

Acceptance level of stink bug egg masses by *T. japonicus* was universally high, ranging from 75% (*C. ligata*) to 100% (*E. conspersus*) on average for each host species, although it was less than 75% for the smallest *P. maculiventris* egg masses and less than 50% for the smallest *C. ligata* egg masses (Figure 3). *Trissolcus japonicus* oviposited in the eggs of all four host species, allocating on average between 0.66 (*C. ligata*) and 1.13 (*E. conspersus*) ovipositions to each available host egg (Figure 3).

### Host survival

The proportion of eggs surviving in egg masses encountered by *T. japonicus* depended on egg mass size only for *C. ligata* (Table 2); for this species, the proportion of surviving eggs followed a quadratic relationship with egg mass size (Figure 3). Survival of *C. ligata* was highest for small egg masses, lowest for intermediate-sized egg masses, and then increased again for the largest egg masses (Figure 3).

The survival of all four host species was reduced (relative to unexposed egg masses) by exposure to *T. japonicus* (Figure 3). The proportion of surviving hosts in egg masses not exposed to parasitoids was not associated with egg mass size (GLMs with quasi-binomial errors; p > 0.05).

### Parasitoid development and host egg abortion

With increasing egg mass size, the number of *T. japonicus* offspring emerging per *H. halys* egg decreased while the proportion of aborted eggs increased (Table 2; Figure 4). For *E. conspersus*, the number of offspring per egg increased with egg mass size but there was no change in the proportion of aborted host eggs. There were no changes in the number of parasitoid offspring per egg or abortion levels with increasing egg mass size for *P. maculiventris* or *C. ligata*.

**Figure 4.**
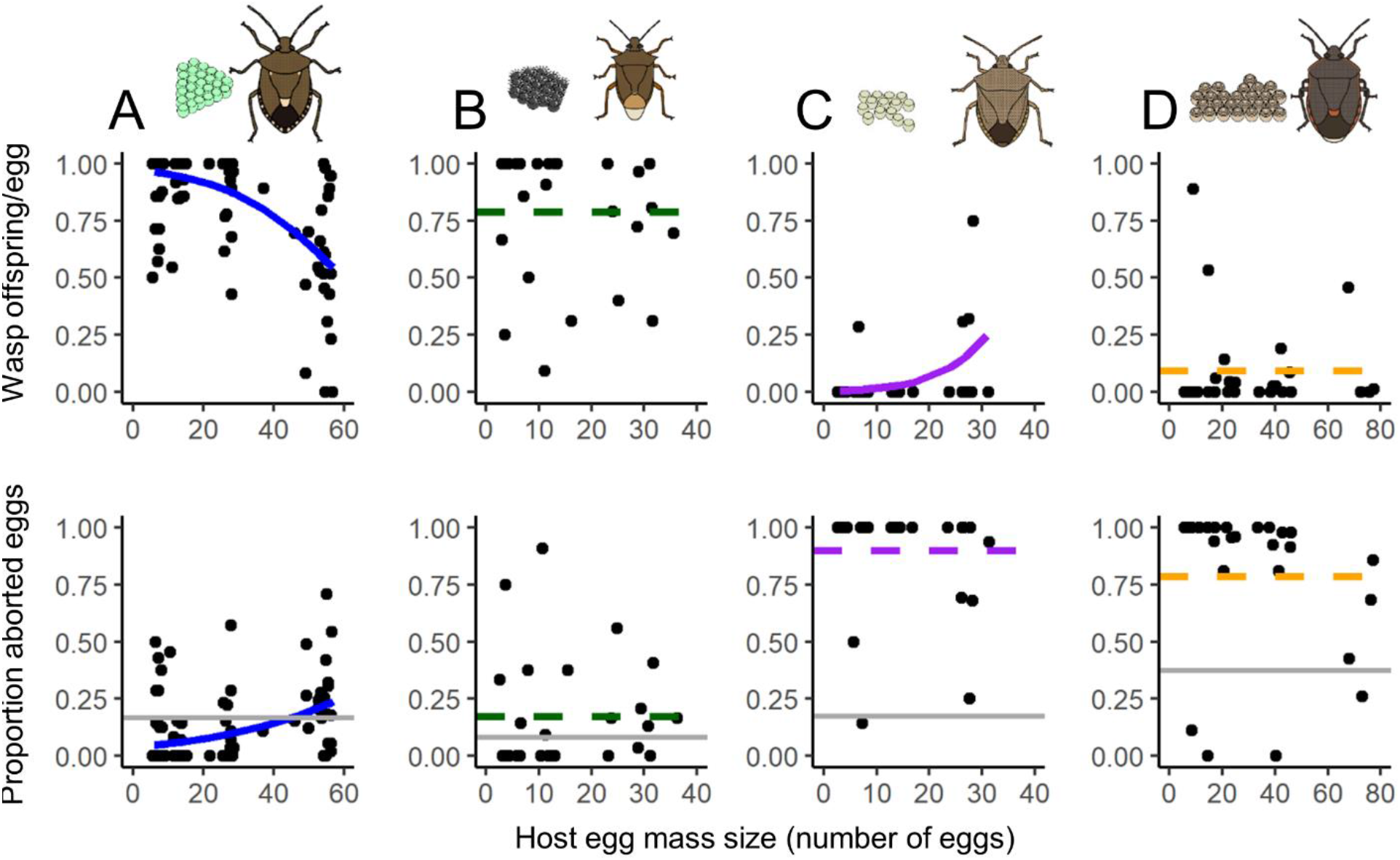
The effect of host egg mass size on the number of *Trissolcus japonicus* wasps emerging per host egg (for egg masses accepted by wasps) and the proportion of aborted host eggs (for egg masses accepted by wasps; solid grey lines show mean host egg abortion in egg masses not exposed to parasitoids) for eggs of four different host species: A) *Halyomorpha halys*; B) *Podisus maculiventris*; C) *Euschistus conspersus*; D) *Chlorochroa ligata*. Solid lines show statistically significant GLM fits; horizontal dashed lines show global means when the relationship between the response variable and host egg mass size was not statistically significant (see Table 2 for statistical information).

*Halyomorpha halys* and *P. maculiventris* were very suitable hosts for *T. japonicus* development; >75% of the eggs of these species, on average, yielded emerging parasitoid offspring and <12% of eggs aborted their development when egg masses were accepted by parasitoids. Despite often being accepted for oviposition by *T. japonicus* (see above), eggs of *E. conspersus* and *C. ligata* rarely (<7% of eggs) yielded emerging parasitoid offspring and had elevated abortion levels compared to egg masses not exposed to parasitoids (Figure 4). Aborted eggs of *C. ligata* in egg masses accepted by *T. japonicus* usually (71% of all aborted eggs pooled among replicates; n = 349) contained fully developed *T. japonicus* adults that failed to emerge from inside host eggs, while aborted eggs of *E. conspersus* usually (97%; n = 726) contained undifferentiated contents without a developed stink bug nymph or parasitoid.

### Additional behavioural parameters

For larger egg masses of *H. halys*, but not the other three host species, *T. japonicus* oviposition rate was slower on larger egg masses (Appendix S1: Table S1; Figure S2). Wasps were more likely to engage in extended guarding behaviour on larger host egg masses, except on *P. maculiventris* eggs where this positive relationship was statistically marginal. Ovipositor rejections by wasps became more likely (as opposed to ovipositions following ovipositor insertion into eggs) over the course of patch exploitation on *H. halys, P. maculiventris*, and *E. conspersus* egg masses, whereas these rejections became less likely on *C. ligata* as patch exploitation sequences progressed.

## Discussion

While we observed some behavioural and reproductive responses to egg mass size by *T. japonicus* on the four host species tested, there was only a significant egg mass size versus survival relationship for one host species, *C. ligata* (Figure 3). In addition, while this was a significant quantitative relationship, it would not result in a large enough refuge to change qualitative host range testing conclusions – *T. japonicus* reduced the survival of all four host species to below 20%, whether or not the full range of egg mass sizes was considered (Figure 5), which would be considered unacceptable levels of attack under current non-target risk assessment scenarios. Thus, our case study provides an example of how intraspecific variation in a potentially protective trait can change conclusions about refuges from parasitism for some non-target species, but not enough to matter in a practical sense for a biological control program.

**Figure 5.**
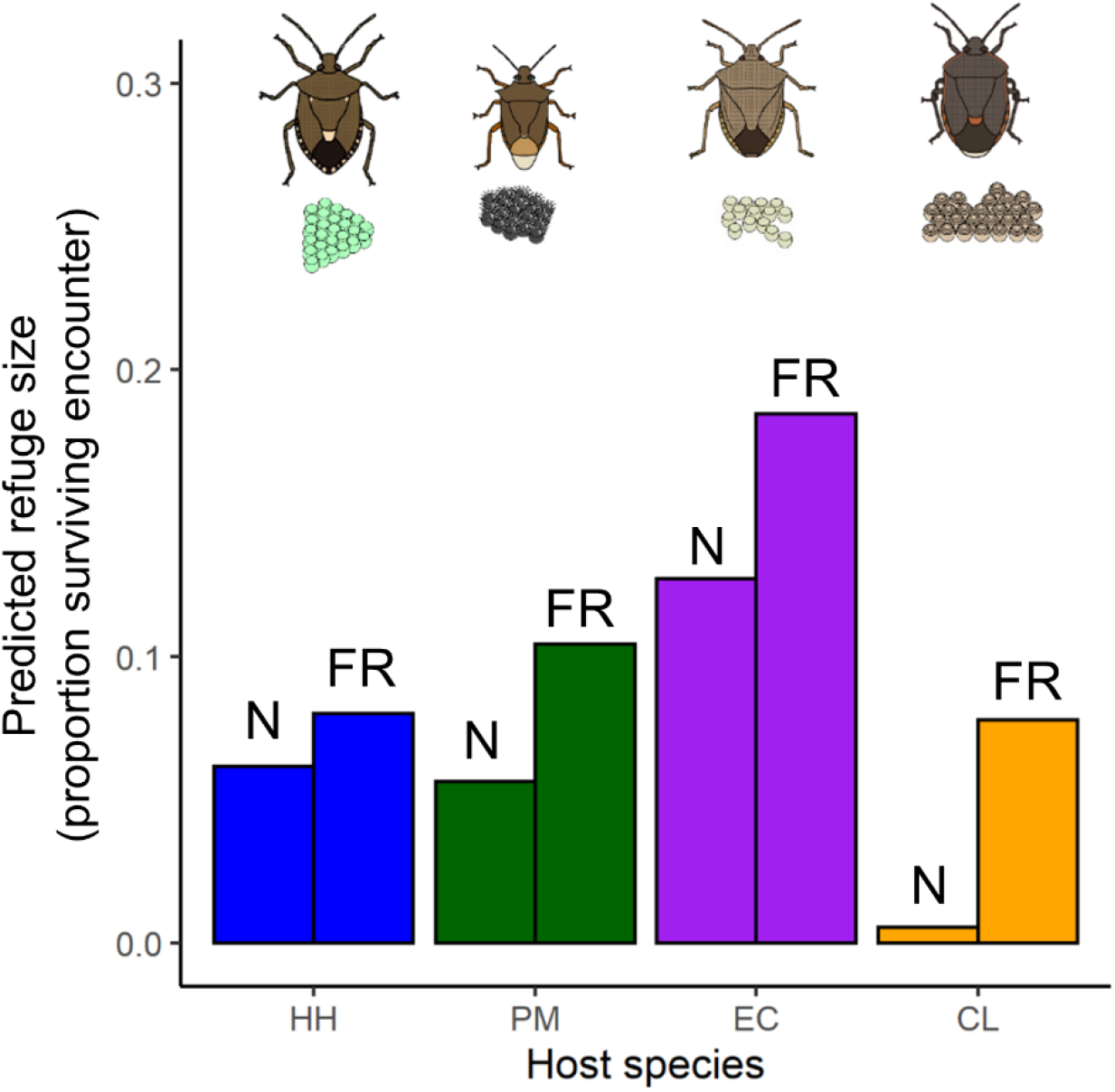
Estimated size of refuges for the four hosts in this study (HH – *Halyomorpha halys*; PM – *Podisus maculiventris*; EC – *Euschistus conspersus*; CL – *Chlorochroa ligata*) either survival of unmanipulated egg masses (N) or the survival of the full range of egg mass sizes (FR) encountered by the parasitoid *Trissolcus japonicus* in our experiments. Refuge size for N was calculated by calculating mean survival for unmanipulated egg masses; refuge size for FR was calculated as outlined in Figure 1; i.e., by multiplying empirically determined *S*(*t*) (bottom row of Figure 3) by *P*(*t*), the proportional frequency of each clutch size for each stink bug species (see Figure 2).

We found mixed evidence for our original predictions that smaller and larger egg masses would have higher survival when exposed to *T. japonicus*. Smaller egg masses were rejected more often by *T. japonicus* only for two of the four host species, *P. maculiventris* and *C. ligata*. Parasitoids showed evidence of egg limitation (i.e., lower numbers of ovipositions per egg) only on host species (*C. ligata, H. halys*) that had artificially manipulated (albeit still within the range of natural variation; see Methods) egg mass sizes that were large enough to deplete their egg loads (∼45-50 eggs; Wong et al. 2021). The combined effects of acceptance probability and egg limitation translated to a significant survival *versus* egg mass size relationship only for *C. ligata*, which displayed a type III susceptibility curve (as per Figure 1). In addition, because actual *C. ligata* egg mass sizes are relatively variable (i.e., small and large egg masses are not extremely rare as they are in *H. halys*, for example; see Figure 2), using the type III relationship to calculate proportion host survival resulted in a ∼14-fold greater estimate of refuge size, although the absolute value of the higher estimated refuge size was still small (<8%; Figure 5). For the other three host species tested, which had type O susceptibility curves, calculating mean survival over the full range of egg mass sizes tested resulted in more modest increases in predicted refuge sizes (Figure 5) as survival was often higher (albeit not significantly so) at lower and/or greater egg mass sizes (Figure 3).

Our results also illustrate that patterns of natural enemy responses to intraspecific variation in host traits may be to some degree host species-specific; qualitative differences in quality among hosts may cause the natural enemy to respond differently to a given trait value depending on the host species. A good example of this from our case study is the fact that *T. japonicus* clearly became egg-limited on large *H. halys* and *C. ligata* egg masses and oviposited fewer times, and this translated to increased host survival in the largest egg masses for *C. ligata* but not *H. halys*. This appeared to be due to *T. japonicus* causing more non-reproductive effects (aborted eggs) on large *H. halys* egg masses (sensu Abram et al. 2016), probably due to the higher frequency of lethal probing during ovipositor rejections when egg-limited on larger *H. halys* (but not *C. ligata*) egg masses (Appendix S1: Figure S2; Table S1). In addition, *T. japonicus* was more likely to reject small egg masses of *C. ligata* (relative to intermediate-sized egg masses), and was more likely to reject individual eggs of *C. ligata* early in its oviposition sequence (Figure S2), but these relationships were not present for *H. halys* eggs. The reasons for these behavioural differences are beyond the scope of the current study, but the result illustrates an important point: intraspecific trait variation may provide a refuge from a given natural enemy in one host species, but not another. Thus, caution should probably be exercised when extrapolating from one host species to others with respect to how quantitative traits determine susceptibility to a natural enemy, especially when testing hosts that have different degrees of shared co-evolutionary history with the natural enemy, as was the case here.

In this study, we manipulated trait values experimentally, which was demanding in terms of host material and made for a more complex experimental design than typical host specificity testing, but had the advantage of providing enough statistical power to determine the response of our natural enemy to less common or less easily obtained host phenotypes. An alternative would be to simply test the range of natural existing trait values, but measure them and include them as a covariate in the study’s analysis. This would have the disadvantage of less often testing rarer trait values and reducing statistical power. Although rarer trait values could be seen as relatively unimportant to the natural enemy’s overall impact on a given host species’ population, they could, for example, be important for preventing a natural enemy’s potential to have large population-level impacts on a given host. For example, if individuals with relatively rare trait values are much less susceptible to natural enemy attacks, it could promote their persistence due to increased selective pressure promoting protected phenotypes (Abrams 2000), even in locations where other phenotypes are heavily attacked. We expected to potentially observe this phenomenon for very small egg mass sizes in our case study system, but this was not the case – based on our laboratory results, small egg mass sizes (even less than 10 eggs) would still be frequently attacked by *T. japonicus*, even though they are much smaller than typical egg masses (28 eggs) laid by their most closely associated host from their native range, *H. halys*. Our results may suggest that for quasi-gregarious parasitoids such as *T. japonicus*, the number of eggs in an egg mass may not be a dominant factor in their assessment of egg mass quality; traits of individual eggs that vary interspecifically, for example, may have relatively more importance (e.g., Sabbatini-Peverieri et al. 2021). Alternatively, as discussed by Murphy et al. (2014), female parasitoids also seek refuges or enemy free space to ensure survival of their offspring, and this may be expressed through variation in certain host-use traits. In the present scenario, the adaptability of *T. japonicus* to exploit a wide range of egg mass sizes may be an important factor in ensuring enemy free space from top-down selective forces (such as predation or hyperparasitism), with the assumption being that smaller egg masses may be less conspicuous and therefore less susceptible to intraguild predation. This would imply a stabilizing trade-off across egg sizes, and could also explain the type O response we observed for most of the pentatomid species tested (Figure 3).

There are a variety of traits that both vary intraspecifically and are associated with differences in susceptibility of hosts to their natural enemies, and that could thus be measured and considered in non-target host range testing. In our case study, we essentially manipulated host size, which is probably an aspect of phenotype that is most obvious and applicable to many host/prey-natural enemy relationships; e.g. because smaller hosts are less suitable and larger hosts may be better defended (Cronin and Strong 1989; McGregor and Roitberg 2000; Heimpel et al. 1996; Abrams 2000). Other traits could include immune defenses such as encapsulation (Kraaijeveld and van Alphen 1999), variable protection conferred by defensive symbionts (Oliver and Higashi 2019), morphological defense traits such as thickness of chorions/integuments, hairs, spines or other physiological traits such as the level of sequestered plant compounds in herbivore host (Gross 1993). In all cases, it is important to ensure that the host phenotypes that are being tested are representative of what occurs in nature. Further, as the frequency distributions of traits are likely to vary among host populations, the individuals tested should ideally be sampled from multiple non-target populations of interest. Trait frequency distributions and trait value-specific susceptibility can only be accurately estimated if sufficient host material is available; if only very small sample sizes of non-target hosts are available (e.g., < 25), testing for candidate trait-based refuges would not be feasible. Finally, any observed relationships between trait values and host susceptibility observed in the laboratory should ideally be followed up in field studies, and could be likewise integrated into post-release evaluation of biological control agents.

Estimating the size of refuges arising from intraspecific variation in protective characteristics is complex and challenging, in part because variation in the behaviour and physiology of natural enemies due to genetic variation and phenotypic plasticity are likely to influence how they respond to intraspecific variation in host traits. For example, if we had tested different geographic populations (e.g., Hopper et al. 2019) or conditions (e.g., ages, nutritional states, experience) (Jenner et al. 2012; Hopper et al. 2013) of *T. japonicus*, they may have responded differently to variation in host egg mass size and may have been more likely to reject certain egg mass sizes or become egg-limited on others (e.g. Hopper et al. 2019). Studying the ecological and evolutionary consequences of interacting intraspecific phenotypic variation in natural enemies and their hosts and prey has a long history in basic evolutionary ecology research (Bolnick et al. 2011; Abrams 2000; Ives and Hochberg 2000), and there is still ample opportunity to apply resulting insights to biological control programs (Roitberg 2000), including refinement of the pre-release predictive capacity of host range testing.

## Acknowledgements

We thank Chris Hou, Sasha Tuttle, Nemo DeJong and Angela Oscienny for help with experiments, dissections, and video analysis; Kennedy Bolstad, Matt Walz, Warren Wong and the Agassiz RDC Greenhouse staff for help with insect rearing. Thanks to George Heimpel, Urs Schaffner, Ryan Paul, Chandra Moffat and Michelle Franklin for helpful discussions and comments on the ideas in this manuscript. This project was supported by Agriculture and Agri-Food Canada, A-BASE projects #2362 and #2955.

## Appendix S1

**Figure S1.**
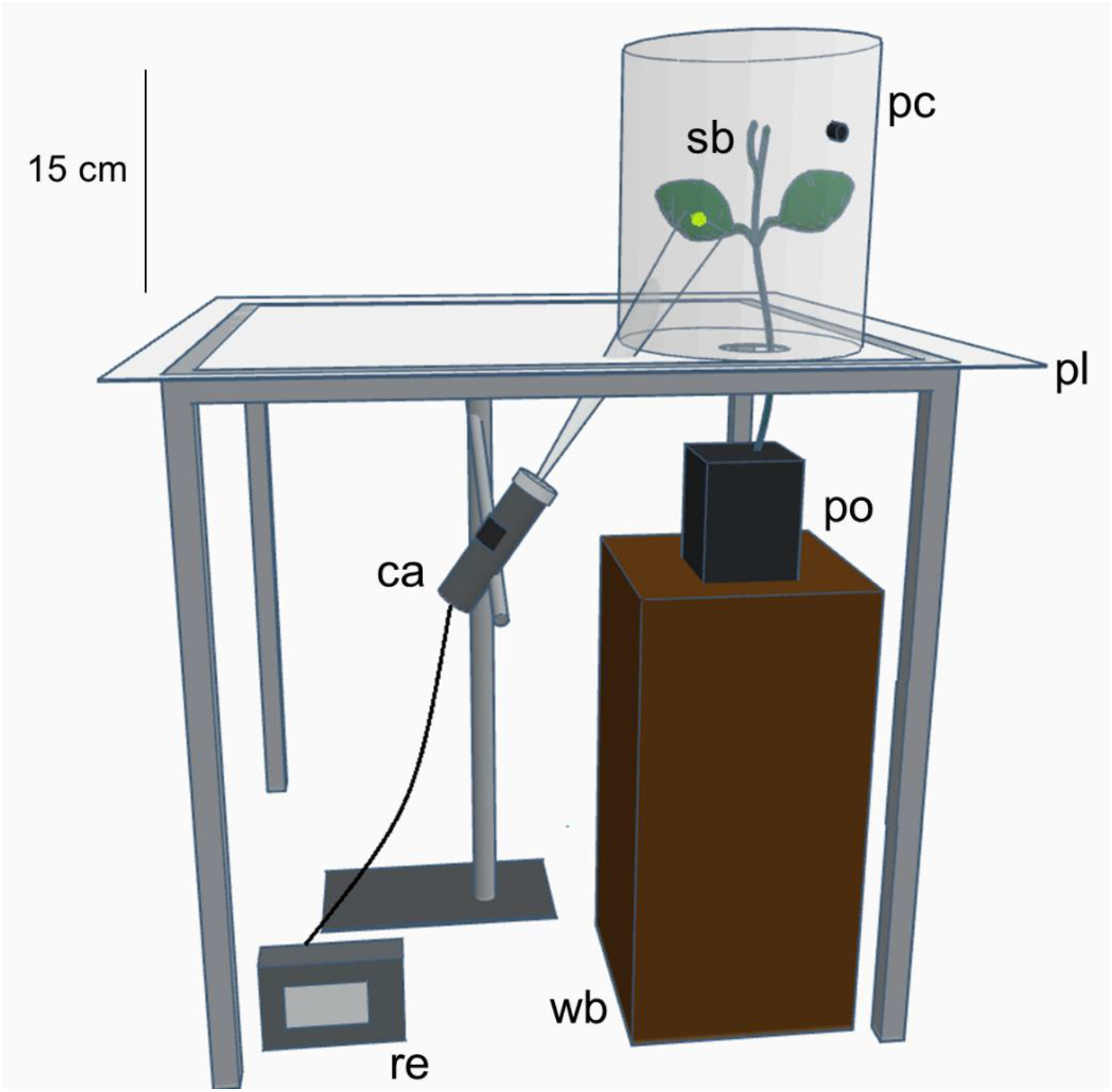
A schematic of the experimental arenas used in this study. pc – transparent plastic cylinder; sb – soybean (*Glycine max* L.) plant; pl – transparent plexiglass sheet; ca – camera; po – plant pot; wb – wooden support block; re – video recording and storage device.

## Methods S1

We also calculated three additional parameters to describe *T. japonicus* behaviour on different host species and egg mass sizes in support the interpretation of the other parameters described in the main text (Figures 3, 4; Table 2). First, we calculated oviposition rate – the number of ovipositions divided by the time between the first and last oviposition, as an index of how efficiently *T. japonicus* exploited host egg masses. Second, because many egg parasitoids in the family Scelionidae, including *T. japonicus*, are known to engage in post-oviposition brood guarding that could affect how long they stay on egg masses of different sizes (Field 1998; Haye et al. 2021), we also recorded the probability of extended brood guarding, defined as whether or not parasitoids remained on the host egg mass for at least 2 hours after the last oviposition, or until the end of the photophase of the first day (as on some larger egg masses, the last oviposition was within 2 hours of the end of the day; similar results were obtained with other criteria for defining extended guarding behaviour). Third, we determined how likely ovipositor rejections versus ovipositions were as parasitoids exploited patches of different hosts and egg mass sizes by scoring the probability of rejections rather than ovipositions (0, 1) over the temporal sequence of patch exploitation for each individual. Statistical methods followed the general procedures outlined in the main text (Methods – ‘Statistical analysis’) and are listed in the footer of Table S1.

**Table S1.** Statistical significance of host egg mass size on behavioral parameters of *Trissolcus japonicus* exploiting the eggs of four different stink bug species, and the change in the probability of rejections over the temporal sequence of patch exploitation. Statistics are bolded when the explanatory variable is statistically significant predictor of the response variable (p < 0.05).

| Response variable | Explanatory variable | Host species |  |  |  |
| --- | --- | --- | --- | --- | --- |
|  |  | *H. halys* | *P. maculiventris* | *E. conspersus* | *C. ligata* |
| Oviposition rate <sup>a</sup> | Host egg mass size | **$F_{1,103} = 29.0$**<br>**$p < 0.0001$** | $F_{1,27} = 0.0$<br>$p = 0.99$ | $F_{1,25} = 0.1$<br>$p = 0.72$ | $F_{1,25} = 0.8$<br>$p = 0.39$ |
| Probability of extended guarding <sup>b</sup> | Host egg mass size | **$\chi^2_{1,103} = 15.4$**<br>**$p < 0.0001$** | $\chi^2_{1,27} = 3.7$<br>$p = 0.053$ | **$\chi^2_{1,25} = 12.8$**<br>**$p = 0.00034$** | **$\chi^2_{1,26} = 7.9$**<br>**$p = 0.0050$** |
| Probability of rejection (vs. oviposition) <sup>c</sup> | Temporal rank of behaviour | **$\chi^2_1 = 5709$**<br>**$p < 0.0001$** | **$\chi^2_1 = 4.6$**<br>**$p = 0.031$** | **$\chi^2_1 = 4.9$**<br>**$p = 0.027$** | **$\chi^2_1 = 22.6$**<br>**$p < 0.0001$** |

<sup>a</sup> GLM with Gaussian error distribution (=linear models)
<sup>b</sup> GLM with binomial error distribution
<sup>c</sup> GLMM with binomial error distribution

**Figure S2.**
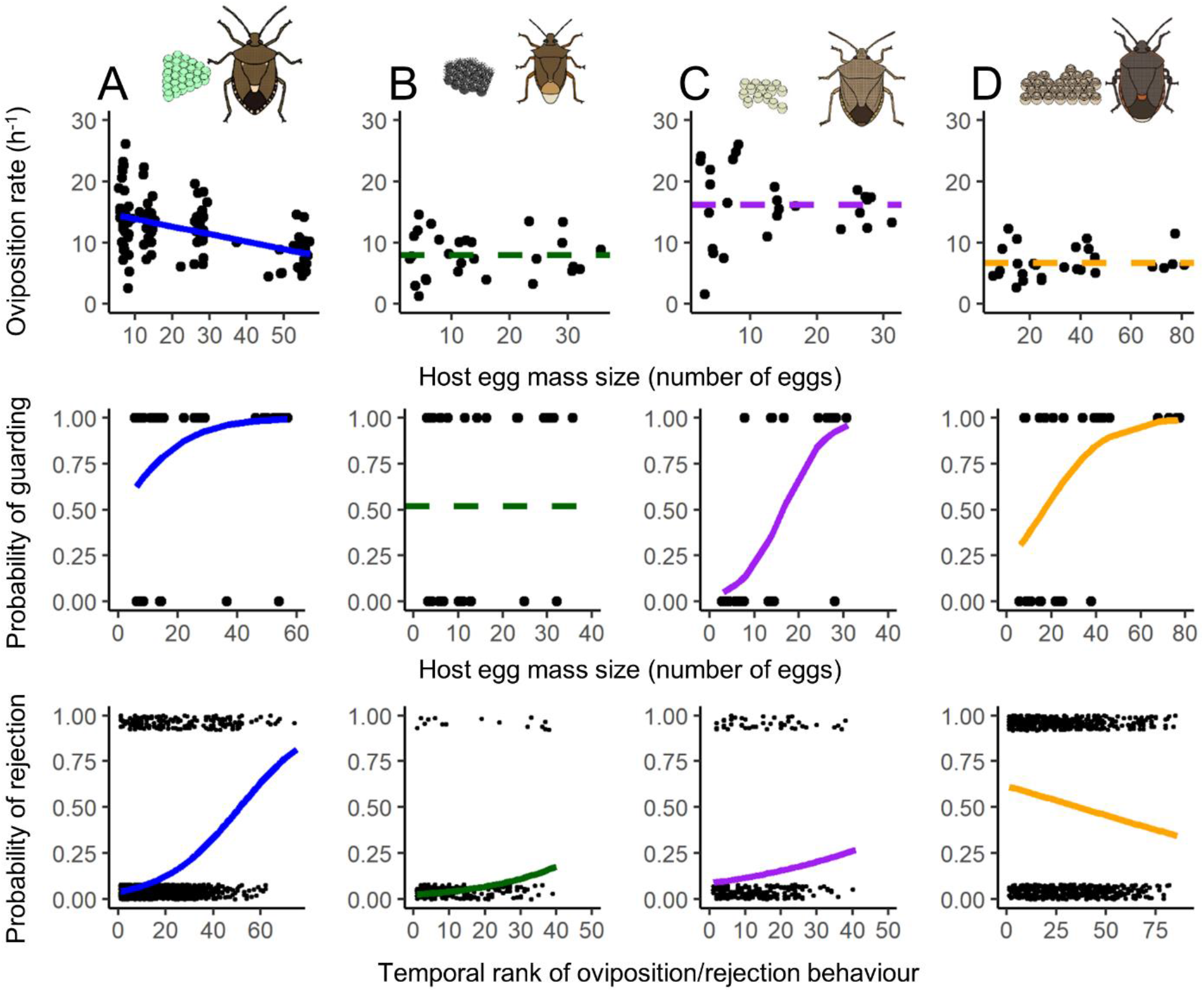
The effect of host egg mass size on *Trissolcus japonicus* patch exploitation behaviour: oviposition rate (the number of ovipositions per hour) and the probability of extended guarding behaviour (probability of remaining on the egg mass for at least 2h or until the end of the trial day), and the probability of rejecting a host egg *versus* ovipositing over time. A) *Halyomorpha halys*; B) *Podisus maculiventris*; C) *Euschistus consperus*; D) *Chlorochroa ligata*. Solid lines show statistically significant GLM (oviposition rate, probability of guarding) or GLMM (probability of rejection) fits; horizontal dashed lines show global means when the relationship between the response variable and host egg mass size was not statistically significant (see Table S1 for statistical information).

